# The growth and pathogenesis of *Citrobacter rodentium* is compromised by disrupted mucin sugar pathways that accumulate N-acetylglucosamine 6-phosphate

**DOI:** 10.1101/2025.08.18.670787

**Authors:** Zhiquan Clarence Huang, Matthias Ahmad Mslati, Caixia Ma, Hyunjung Yang, Qiaochu Liang, Shauna Crowley, Roger Dyer, Irvin Ng, Hongbing Yu, Bruce A. Vallance

## Abstract

Many enteric bacterial pathogens, including the attaching/effacing (A/E) *Escherichia coli* strains, cause acute gastroenteritis in humans. Considering the highly competitive nature of the mammalian gastrointestinal (GI) tract, these pathogens must rely on specific metabolic adaptations to establish successful infections. We hypothesized that A/E pathogens exploit host-derived nutrients within GI mucus, including the monosaccharides N-acetylglucosamine (GlcNAc) and N-acetylneuraminic acid (NeuNAc) to fuel their pathogenesis. Using *Citrobacter rodentium,* a murine-specific A/E pathogen, we disrupted both GlcNAc and NeuNAc catabolism by deleting *nagA*, which encodes the GlcNAc-6-phosphate (GlcNAc-6P) deacetylase that converts GlcNAc-6P into glucosamine-6-phosphate (GlcN-6P). The Δ*nagA* mutant displayed dramatically impaired colonization in C57BL/6J mice and accumulated significant levels of GlcNAc-6P, unlike the Δ*mana* strain, a mutant lacking all GlcNAc and NeuNAc transporters, suggesting that the attenuation was due to sugar-phosphate stress rather than nutrient deprivation alone. Supplementation with glucosamine (GlcN) restored growth, indicating that dysregulated GlcN-6P synthesis, rather than GlcNAc-6P toxicity, underlies the defect. Furthermore, Δ*nagA* exhibited increased susceptibility to several cell wall-dependent stress conditions, in concert with compromised peptidoglycan biosynthesis due to reduced UDP-GlcNAc synthesis. These findings reveal a previously unrecognized metabolic vulnerability in *C. rodentium* and suggest that targeting sugar-phosphate stress responses may provide a new therapeutic strategy against GI bacterial pathogens.

**Importance:** Enteric pathogens like *Citrobacter rodentium* can exploit sugars, including N-acetylglucosamine and N-acetylneuraminic acid, derived from intestinal mucus to grow and infect their hosts. This study shows that disruption of mucin-derived sugar catabolism impairs the fitness of *C. rodentium* in infecting the murine intestine by causing the accumulation of a toxic intermediate of mucin sugar metabolism. Rather than impaired nutrient acquisition, the bacteria are impaired due to the buildup of N-acetylglucosamine-6-phosphate, which depletes substrates for peptidoglycan synthesis. This metabolic bottleneck weakens the bacterial cell wall, making the pathogen more sensitive to environmental stress. These findings identify a conserved metabolic stress response that could be targeted to combat enteric pathogen infections.

## Introduction

The mammalian gastrointestinal tract (GI) contains a large and diverse community of microorganisms that coexist with the host under homeostatic conditions. To maintain GI health, goblet cells, a specialized subset of intestinal epithelial cells (IECs), synthesize and apically secrete mucin, forming a dense and adherent mucus barrier that physically separates luminal microbes from the underlying IECs (1). This mucus barrier is primarily comprised of the gel-forming secreted glycoprotein mucin-2 (Muc2), which represents a large protein backbone with central proline, threonine, and serine-rich domains that undergo dense O-glycosylation in the goblet cell’s Golgi apparatus (2). These O-linked glycans consist of five different monosaccharides: *N*-acetylglucosamine (GlcNAc), *N*-acetylgalactosamine (GalNAc), *N*-acetylneuraminic acid (NeuNAc), galactose, and fucose (3). While mucus serves as a physical barrier that limits microbial access to host tissue, it also forms a nutrient-rich habitat overlying the impermeable barrier layer (4). This niche mucus layer is heavily colonized by an array of commensals, such as *Bacteroides thetaiotaomicron* and *Akkermansia muciniphilas*, that express glycoside hydrolases to cleave mucin glycans and release free sugars that can support the proliferation of these mucolytic bacteria. (5, 6). Notably, these liberated monosaccharides can also be exploited as nutrient sources by nearby enteric pathogens such as *Salmonella enterica serovar* Typhimurium and *Clostridium difficile*, offering them competitive advantages over commensal microbes by supporting their expansion within the niche mucus layer (7).

Recent studies suggest that attaching and effacing (A/E) pathogens, including enterohemorrhagic *Escherichia coli* (EHEC) and enteropathogenic *E. coli* (EPEC), dwell within, and ultimately cross the intestinal mucus layer to establish infection (8). A/E pathogens are major contributors to infantile diarrhea and mortality worldwide (8–10). To colonize their host’s GI tract, A/E pathogens must not only compete with the diverse resident microbiota but also with commensal *E. coli*, which share an overlapping nutrient spectrum (11). Bertin *et al.* showed that the pathogenic EHEC strain EDL933 consumed all five Muc2-derived monosaccharides more efficiently than commensal *E. coli* when exposed to bovine small intestinal contents, demonstrating a competitive metabolic advantage *in vitro* (12). Even so, the precise mechanisms by which pathogens utilize mucin sugars to promote their growth and how such interactions impact infection dynamics *in vivo* are poorly understood.

Direct human studies of EPEC and EHEC are not feasible, and conventional laboratory mice are highly resistant to these clinically important pathogens. Therefore, *Citrobacter rodentium,* a natural A/E murine pathogen, is widely used to model EPEC and EHEC infections in humans (13–15). To establish infection, *C. rodentium* must cross the protective colonic mucus barrier to reach the intestinal epithelium (16, 17). As a member of the *Enterobacteriaceae* family, *C. rodentium* lacks the glycoside hydrolase enzymes used by some commensal microbes to degrade mucin-bound glycans, meaning they are unable to grow on intact mucins (18–20). Recent work by Liang et al. demonstrated that the monosaccharide NeuNAc can promote the growth of *C. rodentium* as well as stimulate the secretion of two autotransporters, Pic and EspC, which enhance *C. rodentium*’s capacity to degrade mucus and adhere to underlying IECs, facilitating its transition from a luminal to a mucosa-associated niche. A *C. rodentium* mutant unable to import NeuNAc showed significantly impaired colonization and caused attenuated pathology in mice, highlighting the important role of NeuNAc utilization in pathogenesis (21, 22). However, NeuNAc is only one monosaccharide component of the Muc2 glycan, and the relative contribution of other abundant sugars, such as GlcNAc, remains poorly defined. Notably, both NeuNAc and GlcNAc are catabolized via pathways that converge onto a shared intermediate, GlcNAc-6-phosphate (GlcNAc-6P), suggesting that perturbations in one pathway may impact the other. This led us to investigate the role of both NeuNAc and GlcNAc in *C. rodentium* colonization, with a focus on how disruption of these metabolic routes may affect pathogen fitness *in vivo*.

In this study, we investigated the mechanisms by which *C. rodentium* exploits free GlcNAc and NeuNAc within the murine colon, and how disruption of its metabolic pathways may impair *C. rodentium*’s fitness and infectivity. To test the role of NeuNAc and GlcNAc catabolism in *C. rodentium*, a mutant strain (Δ*nagA*) was constructed by deleting the *nagA* gene encoding GlcNAc-6P deacetylase. Much like previous studies in *S.* Typhimurium and *Enterobacter hormaechei* that reported severe colonization defects upon deletion of *nagA* (23, 24). *C. rodentium* Δ*nagA* was severely impaired in infecting its murine hosts. However, rather than reflecting an inability to utilize GlcNAc and NeuNAc as nutrients, we determined that the impaired colonization reflected the toxic accumulation of GlcNAc-6P. Our study shows that *nagA* deletion caused a sugar-phosphate stress that compromised cell wall integrity, increasing *C. rodentium’s* susceptibility to antimicrobial and osmotic stress. Supplementation with GlcN, bypassing the reaction catalyzed by NagA, partially restored Δ*nagA* growth, indicating that the stress phenotype arose from the reduction of the downstream product GlcN-6P. These findings reveal a novel mechanism for reducing *C. rodentium* fitness by disrupting sugar catabolism through NagA mutation, which leads to sugar-phosphate toxicity and provides a potential strategy to limit A/E pathogen infection.

## Results

### C. rodentium utilizes specific monosaccharides that are abundant in the colonic mucus

During its infection of the murine colon, *C. rodentium* initially colonizes the outer mucus layer, subsequently traversing the inner mucus barrier layer to infect underlying IECs, highlighting its close association with colonic mucus (16). Previous work by Liang et al. demonstrated that *C. rodentium* can grow on the mucin-derived sugar NeuNAc but not on intact mucin (22). Building on this work, we examined whether *C. rodentium* can utilize other mucin-derived sugars. We cultured wild-type (WT) *C. rodentium* in minimal media supplemented individually with each of the five major monosaccharides found in Muc2 O-glycans. As shown in Figure 1A, *C. rodentium* grew robustly on GlcNAc and NeuNAc and, to a lesser extent, galactose but failed to grow on GalNAc or fucose. These results demonstrate that *C. rodentium* can utilize a subset of mucin-derived sugars for growth, particularly GlcNAc and NeuNAc, which appear to be preferred carbon sources.

**Figure 1.**
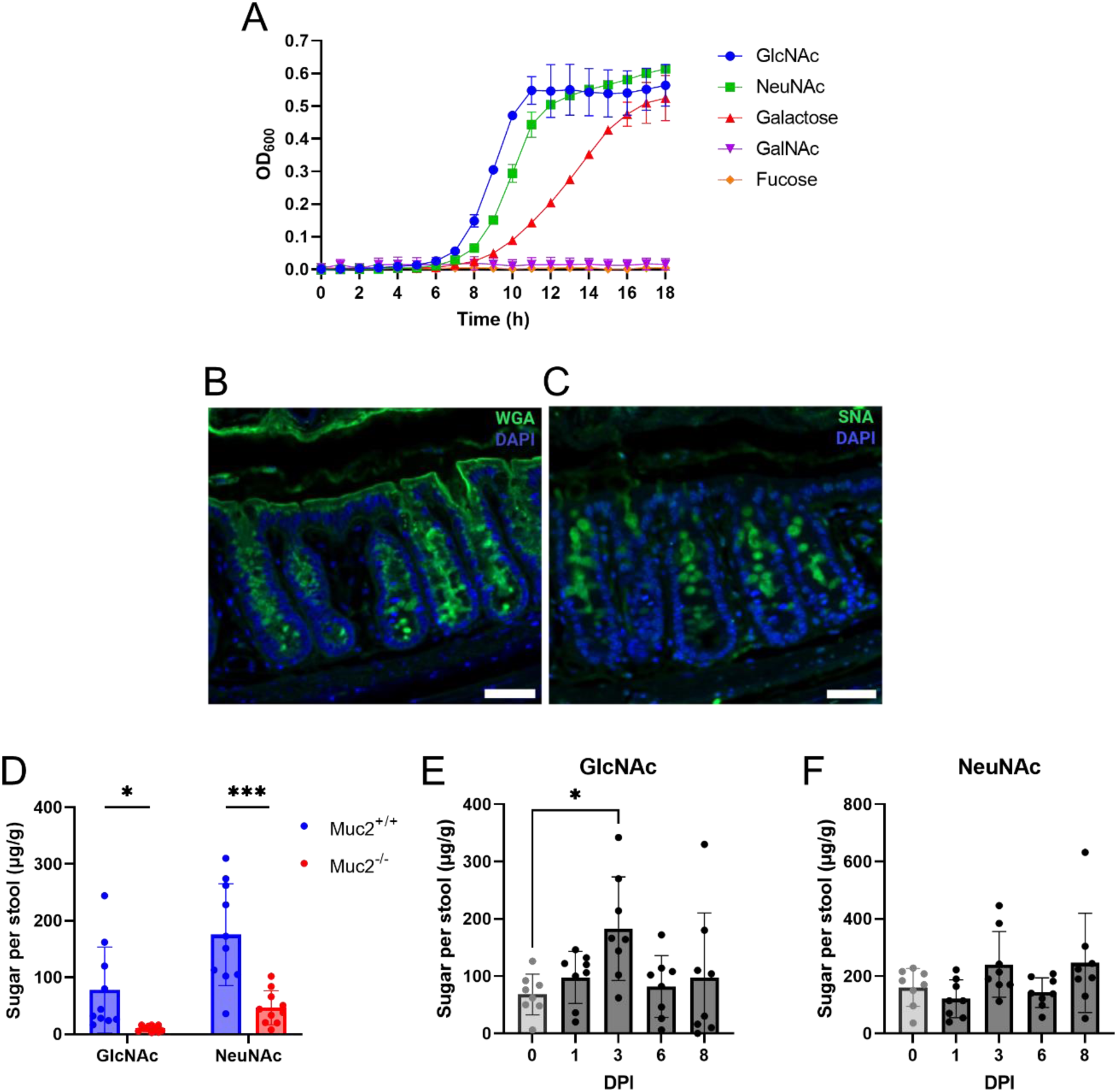
*C. rodentium* utilizes GlcNAc and NeuNAc that are enriched in the murine colon of C57BL/6J mice. A) Growth assay of WT *C. rodentium* in minimal media with five monosaccharides constituting the Muc2 O-glycans. The absorbance at OD_600_ was measured every hour and were shown as Mean ± SD from biological triplicates. B) WGA and C) SNA staining of colonic cross sections from uninfected mice (at baseline). Sections were stained with WGA or SNA (green) to detect GlcNAc or NeuNAc, respectively, and DAPI (blue) to detect DNA. Original magnification, 200X. Images are representative of 3 independent experiments with 5 mice per experiment. (Scale bar, 20 μm.) D) Levels of GlcNAc and NeuNAc in the stools of *Muc2^+/+^* and *Muc2^-/-^* littermates assessed by UHPLC/QqQ-MS. Data were collected from 10 mice pooled from 2 independent experiments and shown in Mean ± SD. Statistical significance was determined by multiple t-test. **P < 0.01. Levels of E) GlcNAc and F) NeuNAc in the stool samples of mice at baseline and during infection with *C. rodentium* assessed by UHPLC/QqQ-MS. Data were collected from 8 mice pooled from 3 independent experiments and shown in Mean ± SD. Y-axis represents the amount of sugar in microgram per gram of stool (D-F). Statistical significance was determined by One-way ANOVA (E). *P < 0.05.

We next tested whether GlcNAc and NeuNAc are present and accessible to *C. rodentium* within the murine colon. For these studies, we chose C57BL/6J mice, as they have been shown to be more susceptible to *C. rodentium* infection than other C57BL/6 substrains (25). To examine the spatial distribution of GlcNAc and NeuNAc, we stained colon tissue sections from C57BL/6J mice with the fluorescently labelled lectins, Wheat Germ Agglutinin (WGA), which preferentially binds to GlcNAc, and Sambucus Nigra Agglutinin (SNA), which has an affinity for α-2,6-linked NeuNAc (26). WGA staining revealed a widespread signal within the cytoplasm of goblet cells as well as strong labelling throughout the mucus layer lining the epithelial surface (Figure 1B). In contrast, SNA staining appeared more punctate and enriched at the apical surface of goblet cells and in the outer mucus layer. (Figure 1C). While both lectins labelled goblet cells and the overlying mucus, their distinct patterns suggest differential localization of GlcNAc and NeuNAc within the mucus barrier, consistent with previous reports using WGA and SNA staining in the mouse colon tissues (22, 27).

To more precisely quantify GlcNAc and NeuNAc, we measured their levels within mouse fecal samples using an ultrahigh-performance liquid chromatography coupled with triple quadrupole mass spectrometry (UHPLC/QqQ-MS). Because these sugars can originate from various sources, including exogenous food and host glycoproteins, we sought to determine whether they are primarily derived from secreted mucins. To do this, we compared wild-type *(Muc2⁺/⁺)* mice with *Muc2^⁻/⁻^* mice, which lack the major gel-forming mucin, Muc2, in the colon. Feces collected from *Muc2^+/+^*mice exhibited significantly higher levels of GlcNAc and NeuNAc as compared to *Muc2^-/-^* mice, supporting the conclusion that these sugars predominantly originate from Muc2 (Figure 1D).

We next assessed whether *C. rodentium* infection altered the abundance of these sugars. Sugar quantification of fecal samples from infected C57BL/6J mice revealed that GlcNAc and NeuNAc remained detectable throughout the infection course (Figures 1E and 1F), indicating their availability as potential nutrients. GlcNAc levels significantly increased by 3 days post-infection (DPI) compared to baseline, suggesting that *C. rodentium* infection may enhance the release of GlcNAc from mucin glycans. Together, these findings confirm that GlcNAc and NeuNAc are spatially enriched in the mucus layer, persist throughout infection, and are available for metabolic exploitation by *C. rodentium*.

### Inactivation of the GlcNAc and NeuNAc catabolic pathway impairs C. rodentium colonization in mice

The catabolic pathways for NeuNAc and GlcNAc are intricately linked among diverse microorganisms, with NeuNAc breakdown feeding directly into the GlcNAc pathway (28). Specifically, NeuNAc is first cleaved into N-acetylmannosamine and then converted into GlcNAc-6P, the same intermediate produced when extracellular GlcNAc is absorbed via the phosphotransferase system (PTS). In the bacterial cytoplasm, NagA further deacetylates GlcNAc-6P to form glucosamine-6-phosphate (GlcN-6P), a precursor for glycolysis and peptidoglycan synthesis (Figure 2A).

**Figure 2.**
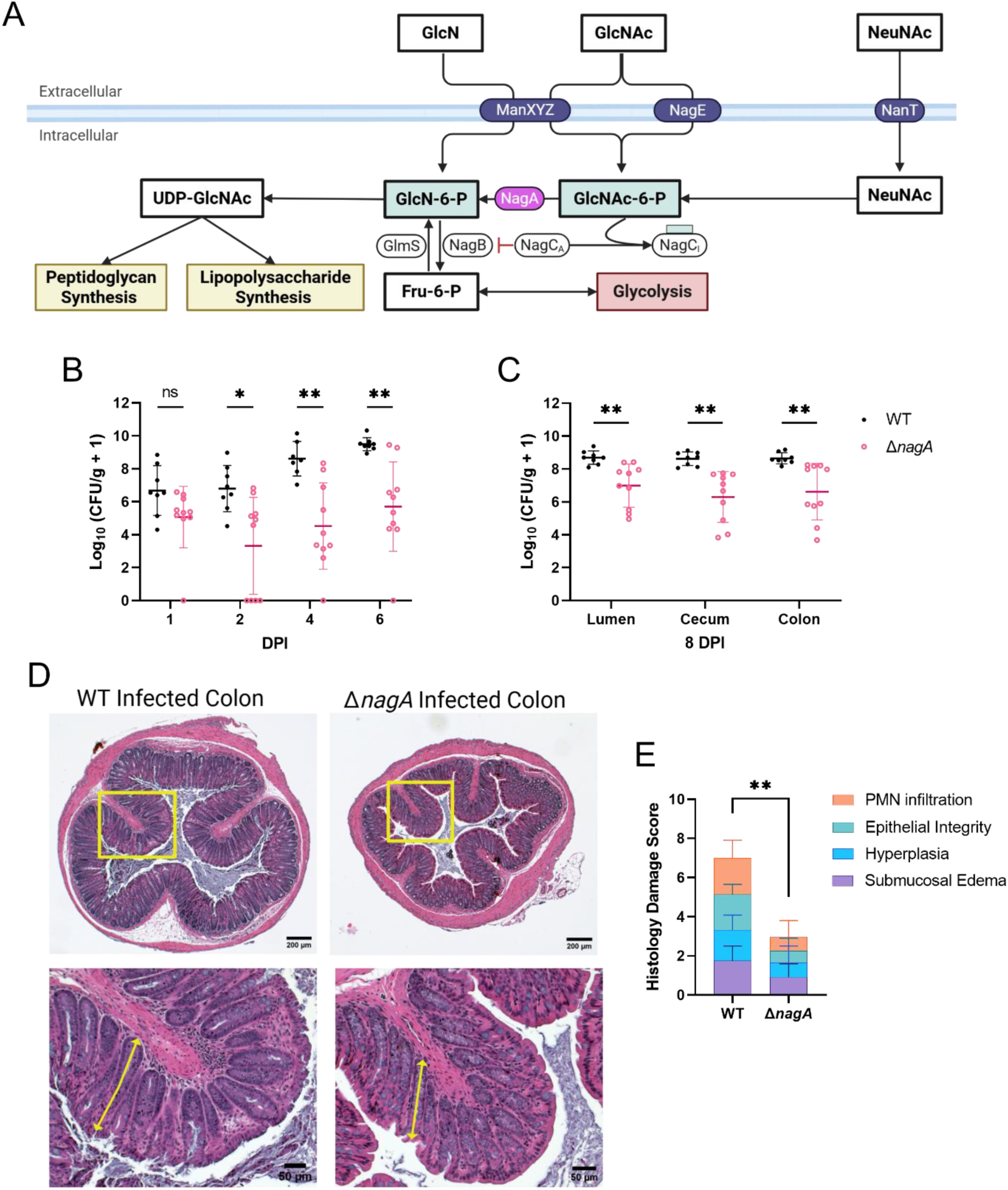
*C. rodentium* Δ*nagA* is significantly impaired in colonizing C57BL/6J mice intestine. A) Schematic representation of the metabolic pathway of GlcNAc and NeuNAc in *C. rodentium*. NeuNAc is transported into the cell via the NanT transporter and converted into GlcNAc-6P, while GlcNAc are taken up by the ManXYZ and NagE PTS to directly form GlcNAc-6P. Inside the cell, GlcNAc-6P is deacetylated by NagA to generate GlcN-6P for glycolysis or peptidoglycan and lipopolysaccharide synthesis. B) *C. rodentium* CFU from stool pellets collected at 1, 2, 4, and 6 DPI and C) luminal contents as well as intestinal tissues collected at 8 DPI from C57BL/6J mice orally infected with 10^8^ CFU of WT (n = 8) or Δ*nagA* (n = 10) *C. rodentium* (n, number of biological replicates). D) Representative H&E-stained distal colon at 8 DPI for WT and Δ*nagA* mutant-infected mice. (Scale bar, 200 μm.) Lower panels are expanded images of corresponding boxed regions in panels above. Crypt hyperplasia is indicated by arrows. (Scale bar, 50 μm.). E) Blinded histopathological scores of H&E tissue sections of mice infected with WT (n = 8) or Δ*nagA* (n = 10) *C. rodentium* (see Methods for scoring criteria). Agreement among raters ensured by Spearman’s rank correlation coefficient r = 0.9283. Data were shown in Mean ± SD and statistical significance was determined by multiple t-tests. ***P < 0.001, **P < 0.01, *P < 0.05 (B, C, E).

To examine the role of GlcNAc and NeuNAc catabolism in *C. rodentium* colonization, we generated a mutant strain, Δ*nagA,* which lacks the NagA enzyme required to convert GlcNAc-6P into GlcN-6P. We infected C57BL/6J mice with either the WT or the Δ*nagA C. rodentium* strain and monitored pathogen shedding in fecal pellets at 1, 2, 4, and 6 days post infection (DPI), followed by cecal and colonic tissue collection at 8 DPI. WT and Δ*nagA* infected mice exhibited similar low pathogen burdens at 1 and 2 DPI, harboring approximately 10^6^ colony-forming units (CFU) per gram of stool (Figure 2B). As the infection progressed, WT displayed a robust expansion to 10^9^ CFU by 6 DPI, while Δ*nagA* remained at levels 100-1000 fold lower. This colonization difference persisted until 8 DPI, where WT *C. rodentium* exhibited significantly higher CFUs in the stool as well as cecal and colonic tissues as compared with the Δ*nagA* mutant strain (Figure 2C).

Histological analysis revealed that the Δ*nagA*-infected mice developed a milder inflammatory response than WT-infected mice (Figure 2D-E). While WT infection led to extensive epithelial damage, immune cell infiltration, and crypt hyperplasia, infection by the Δ*nagA* strain resulted in only limited epithelial cell disruption and reduced immune infiltration. These results demonstrate that the Δ*nagA* strain is impaired in its ability to colonize and induce pathological tissue damage in the colon.

### The attenuated virulence of ΔnagA is independent of GlcNAc/NeuNAc metabolism

As noted, previous studies have reported severe colonization defects of *S.* Typhimurium and *E. hormaechei* upon deletion of *nagA,* which were attributed to their inability to access GlcNAc and NeuNAc as nutrients (23, 24). To explore if this is the basis for the colonization defect observed with Δ*nagA C. rodentium*, we generated a mutant strain that cannot transport GlcNAc and NeuNAc into the cytoplasm. This mutant, lacking ManXYZ, NagE, and NanT transporters, was designated Δ*mana.* To rule out an unintentional polar effect created by the *nagA* deletion, we also generated a Δ*nagA/*pNagA complementation strain where the Δ*nagA* mutant was transformed with a plasmid expressing NagA under its native promoter. WT, Δ*nagA,* Δ*nagA*/pNagA, and Δ*mana* were grown in minimal media supplemented with both GlcNAc and NeuNAc as carbon sources. WT and the Δ*nagA/*pNagA strain demonstrated similar growth rates, while Δ*mana* and Δ*nagA* failed to proliferate (Figure 3A).

**Figure 3.**
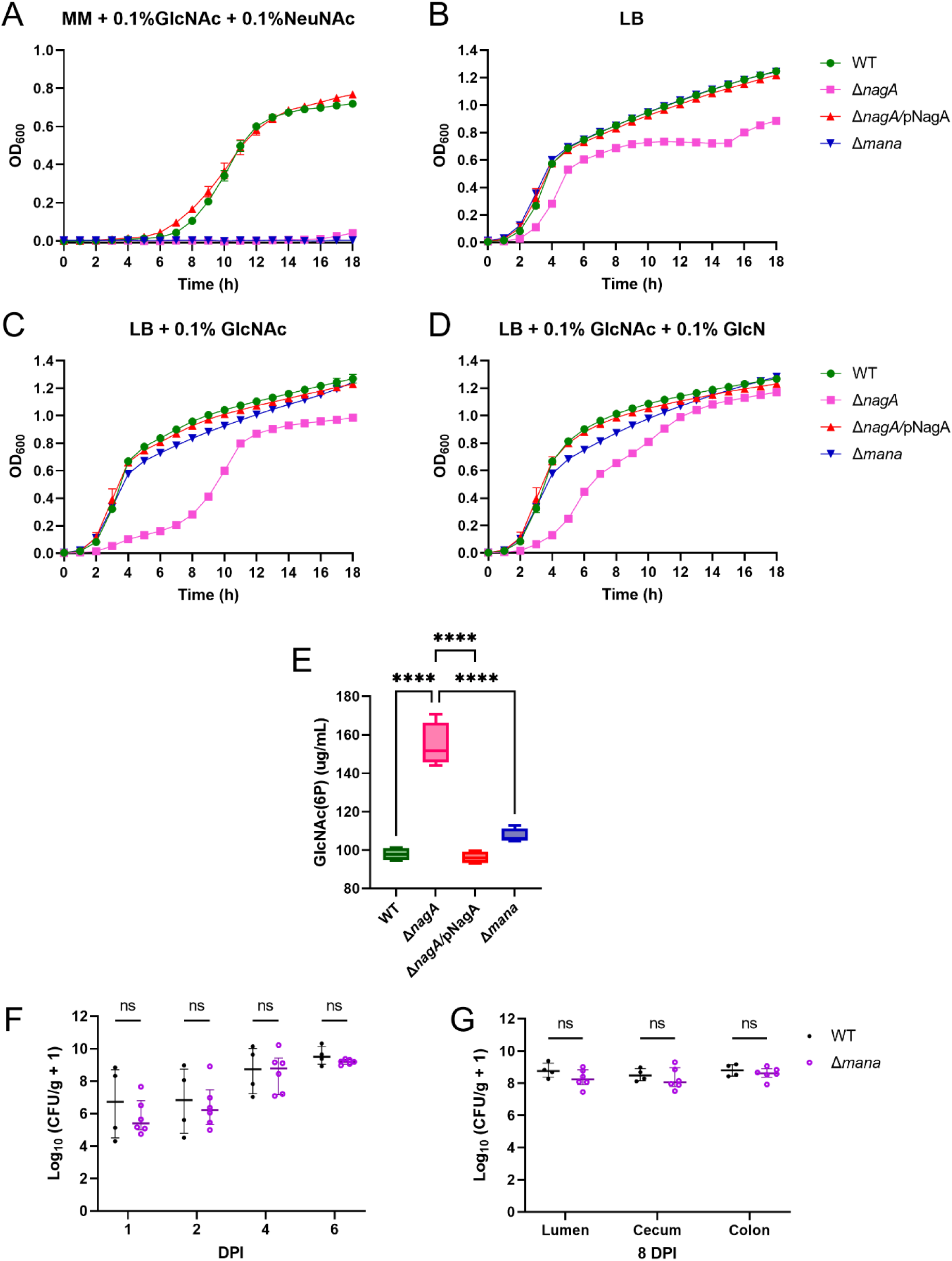
**GlcNAc-6P accumulation leads to growth and colonization defect in Δ*nagA.*** Growth assay of *C. rodentium* WT, Δ*nagA*, Δ*nagA/*pNagA, and Δ*mana* grown in A) minimal medium supplemented with 0.1% GlcNAc and NeuNAc, and B) LB media alone, C) or supplemented with 0.1% GlcNAc and D) 0.1 % GlcN. The absorbance at OD_600_ was measured every hour shown as Mean ± SD from biological triplicates. E) The levels of GlcNAc(6P) in the soluble extracts of the four *C. rodentium* strains were measured by the modified Morgan-Elson assay. Data was represented as Mean ± SD from biological quadruplicates. Statistical significance was determined by One-way ANOVA. ****P < 0.0001. C57BL/6J mice were orally infected with 10^8^ CFU of WT (*n* = 4) or Δ*mana* (*n* = 6) (*n*, number of biological replicates) *C. rodentium*, and the CFU was enumerated from F) stool pellets collected at 1, 2, 4, and 6 DPI; G) luminal contents as well as intestinal tissues collected at 8 DPI. Data was shown in Mean ± SD and statistical significance was determined by multiple t-tests (F-G).

To examine whether the metabolic defect of Δ*nagA* could be rescued in nutrient-rich conditions containing abundant alternative carbon sources, such as glucose and amino sugars, we cultured all strains in LB media. Surprisingly, Δ*nagA* exhibited a distinct growth delay and failed to reach the same final optical density as the WT, Δ*mana*, or Δ*nagA/*pNagA strains (Figure 3B). This growth defect was further exacerbated by supplementation with GlcNAc (Figure 3C), but, interestingly, it was partially rescued by supplementation with glucosamine (GlcN), which is a carbon source that can bypass the need for NagA (Figure 3D). Analysis of the growth curves indicates that the growth defect of Δ*nagA* in nutrient-rich conditions is not solely driven by the inability to catabolize GlcNAc or NeuNAc as carbon sources, but likely due to the introduction of a chokepoint within the GlcNAc/NeuNAc catabolic pathway.

Although both Δ*nagA* and Δ*mana* interrupt GlcNAc and NeuNAc catabolism, they disrupt this pathway at different points. Δ*nagA* blocks the conversion of the GlcNAc- and NeuNAc-derived catabolic product, GlcNAc-6P, to GlcN-6P. However, it does not interrupt the transport of GlcNAc via the PTS nor the initial steps of NeuNAc catabolism. As a result, both sugars can continue to be imported and catabolized, but are unable to proceed past the formation of GlcNAc-6P. In contrast, the Δ*mana* strain lacks these transporters and is unable to import either sugar (Figure 2A). In *Enterobacteriaceae* such as *C. rodentium* and *E. coli*, ManXYZ is a PTS transporter with broad specificity capable of importing glucose, mannose, GlcNAc and GlcN into the cytoplasm as phosphorylated intermediates (e.g., GlcNAc-6P, GlcN-6P) (29). Unlike GlcNAc, which requires NagA to be converted to GlcN-6P, GlcN enters the pathway as GlcN-6P directly and bypasses the blocked step in the Δ*nagA* strain. Therefore, we hypothesized that the attenuated phenotype of Δ*nagA* resulted from GlcNAc-6P accumulation and GlcN-6P depletion.

We next quantified intracellular GlcNAc-6P levels in the WT, Δ*nagA*, Δ*mana*, and Δ*nagA/*pNagA *C. rodentium* strains using the Morgan-Elson assay. This colorimetric assay detects N-acetylhexosamine (including GlcNAc and GlcNAc-6P) using alkaline borate, followed by Ehrlich’s reagent to produce a chromogenic product measurable at 585 nm (30). When our strains were grown in minimal media supplemented with glucose and GlcNAc, the cytosolic level of GlcNAc-6P in the Δ*nagA* strain was found to be significantly higher than WT, Δ*nagA/*pNagA complemented strain, and Δ*mana* (Figure 3E). These data demonstrate that Δ*nagA* accumulates GlcNAc-6P intracellularly.

To clarify whether GlcNAc and NeuNAc uptake contribute to the colonization of *C. rodentium* in the murine intestine, we infected C57BL/6J mice with WT or Δ*mana*. In contrast to Δ*nagA,* which showed a marked colonization defect (Figure 3G and 3H), Δ*mana*-infected mice carried high pathogen burdens comparable to mice infected with WT. Interestingly, in a competitive infection assay, WT outcompeted Δ*mana,* demonstrating that GlcNAc/NeuNAc catabolism does provide a fitness advantage to *C. rodentium* during initial colonization but not to the level seen with Δ*nagA* (Supplementary Figure 1). The above results indicate that the attenuated phenotype of the Δ*nagA* mutant cannot be solely explained by its inability to catabolize GlcNAc/NeuNAc but instead suggests that, through the deletion of *nagA*, a metabolic chokepoint is created, leading to GlcNAc-6P accumulation and compromised interconnected metabolic pathways, overall presenting as attenuated virulence *in vivo*.

### GlcNAc-6P accumulation disrupts normal GlcN-6P synthesis in ΔnagA, leading to compromised cell wall integrity

Following the observation that GlcNAc-6P accumulated in the Δ*nagA* mutant and potentially impaired its growth and *in vivo* colonization, we next explored possible mechanisms driving this phenotype. We incubated WT, Δ*nagA*, Δ*mana*, and Δ*nagA/*pNagA *C. rodentium* strains in PBS supplemented with 0.1% GlcNAc for 18 hours and enumerated the CFU before and after incubation to explore if the accumulation of GlcNAc-6P produced a cytotoxic effect. Consistent with the previous growth assay in minimal media, WT and the complemented Δ*nagA*/pNagA strains multiplied, while the Δ*nagA* and Δ*mana* mutants did not. Importantly, similar to Δ*mana*, no significant decline in viability was observed in Δ*nagA*, suggesting the accumulation of GlcNAc-6P in Δ*nagA* does not directly kill *C. rodentium* (Figure 4A).

**Figure 4.**
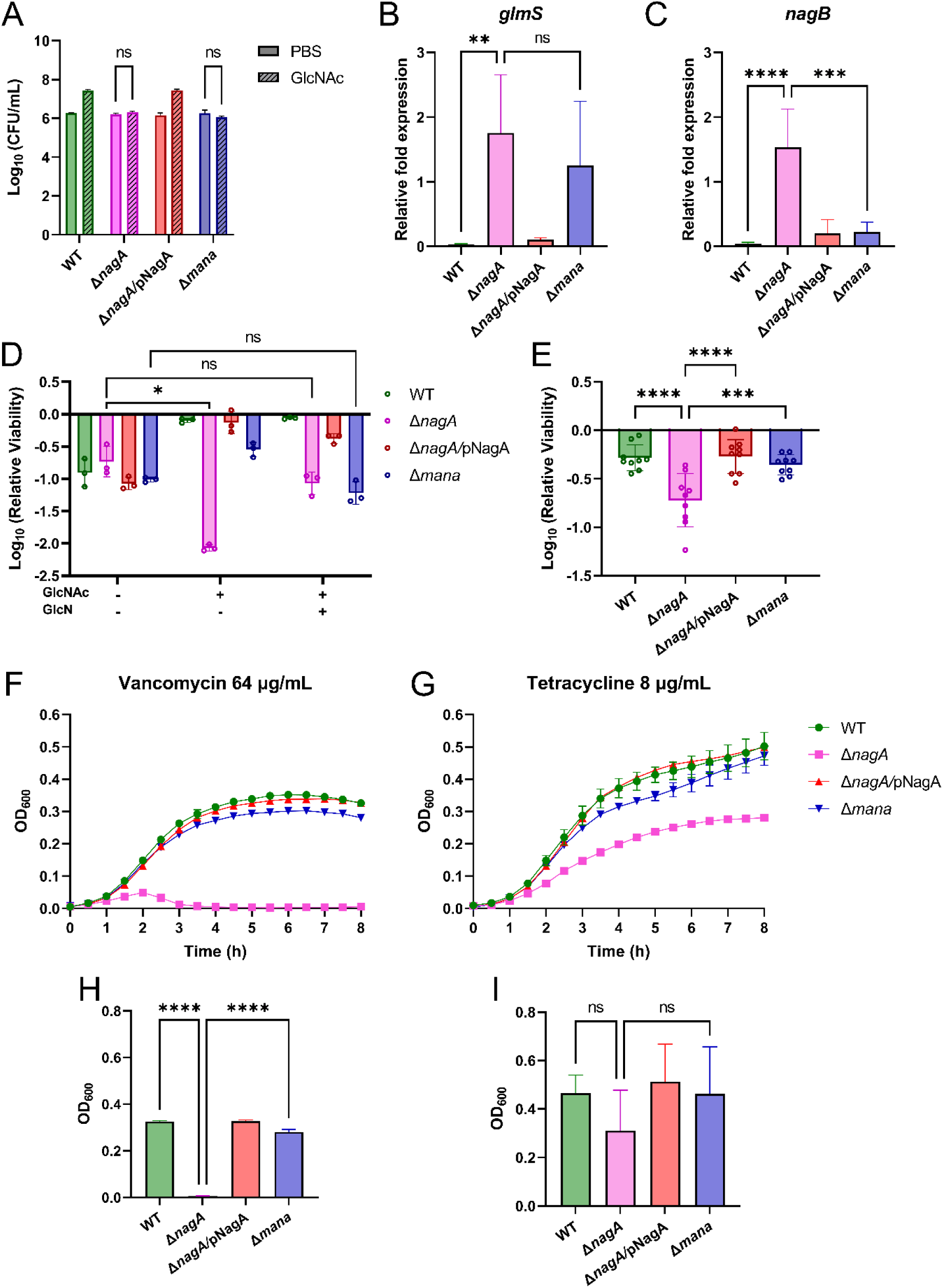
GlcNAc-6P accumulation depletes GlcN-6P that leads to increasing sensitivity to cell envelop stressors. A) Bacterial concentration of WT, Δ*nagA*, Δ*nagA/*pNagA, and Δ*mana C. rodentium* grown in PBS supplemented with or without 0.1% GlcNAc for 24h. Statistical significance was determined by multiple t test. Bacterial qPCR analysis of genes responsible for B) GlcN-6P synthesis from Fru-6P (*glmS*) and C) conversion to Fru-6P (*nagB*). The expression level was normalized to the relative expression of the reference gene, *rpoA*, using the 2^-ΔCᴛ^. D) Survival of *C. rodentium* in the presence of 10mg/mL lysozyme supplemented with/without 0.1% GlcNAc and GlcN. Relatvie viability was assessed by CFU quantification, normalized to PBS control groups. E) Survival of *C. rodentium* in LB under osmotic stress induced by 3% NaCl. Relative viability was assessed by CFU quantification, normalized to 1% NaCl control groups. Growth curve of *C. rodentium* in LB supplemented with F) 64 μg/mL vancomycin or H) 8 μg/mL tetracycline. OD_600_ was monitored every half hour for a total of 8 hours. The OD_600_ at the 8 h of WT, Δ*nagA*, Δ*nagA/*pNagA, and Δ*mana C. rodentium* after 8 hours incubation in G) vancomycin and I) tetracycline from three independent experiments. All the data were shown in Mean ± SD. Statistical significance was determined by One-way ANOVA (B-E, H-I). ****P < 0.0001, ***P < 0.001, **P < 0.01, *P < 0.05.

Next, the impact of accumulated GlcNAc-6P accumulation on gene expression within the GlcNAc and NeuNAc catabolic pathway was evaluated, where we measured the transcript levels of *glmS* and *nagB* using RT-qPCR. *glmS* encodes an aminotransferase that synthesizes GlcN-6P from fructose-6-phosphate (Fru-6P), while *nagB* encodes a deaminase that converts GlcN-6P to Fru-6P (Figure 2A). WT and Δ*nagA/*pNagA, which are able to generate GlcN-6P from exogenous GlcNAc, showed significantly lower *glmS* expression levels compared with Δ*nagA* and Δ*mana*, confirming that the latter strains generate GlcN-6P from endogenous Fru-6P (Figure 4B). However, it was observed that Δ*nagA* also significantly upregulates *nagB* compared with the other three strains (Figure 4C). Previous studies in *E. coli* have shown that GlcNAc-6P accumulation deactivates the *nag* operon repressor, NagC, resulting in the derepression of *nagA* and *nagB* (31, 32). Consistent with the findings in *E. coli*, these results highlight a regulatory dilemma for Δ*nagA*: GlcNAc-6P accumulates but cannot be deacetylated to GlcN-6P, leaving *nagB* constitutively active to impede GlcN-6P synthesis. In contrast, Δ*mana* can synthesize GlcN-6P from Fru-6P and does not need to combat NagB upregulation, as was observed in the *ΔnagA* strain due to GlcNAc-6P accumulation, thus avoiding this regulatory and metabolic imbalance. Taken together, these results demonstrate that GlcNAc-6P accumulation in the absence of *nagA* leads to the upregulation of *glmS* and *nagB* gene expression.

GlcN-6P is the essential substrate for peptidoglycan and lipopolysaccharide synthesis. We next assessed whether the compromised GlcN-6P synthesis could affect the integrity of the *C. rodentium* cell wall and membrane. Lysozyme is an innate antimicrobial enzyme found in the mammalian GI tract that hydrolyzes the β-1,4-glycosidic bond in the peptidoglycan backbone, thereby compromising bacterial cell wall integrity (33). To assess whether GlcNAc-6P accumulation sensitizes *C. rodentium* to lysozyme, we exposed our *C. rodentium* strains to 10 mg/mL lysozyme in the presence of GlcNAc or GlcN. It was observed that Δ*nagA* was more sensitive to the lysozyme challenge in the presence of GlcNAc, and such sensitivity was reduced when GlcN was added (Figure 4D). This is consistent with our previous finding that GlcN bypasses the blocked NagA-dependent step by entering the pathway as GlcN-6P, thereby alleviating the negative effect of GlcNAc-6P accumulation and restoring cell wall synthesis.

Moreover, Δ*nagA* exhibited decreased viability under high NaCl concentrations (Figure 4E), indicating higher sensitivity to osmotic stress. Antibiotic susceptibility profiling further supported this envelope defect: Δ*nagA* proved more vulnerable to vancomycin, a glycopeptide targeting bacterial cell wall through blocking cross-linking by binding D-Ala-D-Ala termini of peptidoglycan precursors (Figure 4F and 4H). In contrast, Δ*nagA*’s susceptibility to tetracycline, a ribosome-targeting antibiotic that inhibits protein synthesis without directly affecting peptidoglycan, remained similar to the other three strains. (Figure 4G and 4I). These results collectively demonstrate that GlcNAc-6P accumulation disrupts cell wall integrity through depleting GlcN-6P pools, providing a mechanistic explanation for the attenuated colonization observed in the *C. rodentium* Δ*nagA* strain. Altogether, GlcNAc-6P accumulation impairs GlcN-6P biosynthesis, disrupting peptidoglycan homeostasis and sensitizing Δ*nagA* to cell wall stressors, likely contributing to its impaired *in vivo* colonization.

## Discussion

Our findings demonstrate that disruption of the sugar metabolism enzyme NagA impairs *C. rodentium*’s colonization and ability to cause colonic pathology in infected mice. Notably, several recent studies have identified similar major virulence defects in *S.* Typhimurium and *E. hormaechei* (23, 24). Strikingly, we determined that this defect in pathogenesis was not due to an inability to use the mucin-derived sugars GlcNAc and NeuNAc as nutrients, but instead reflects the creation of a metabolic bottleneck, driving the accumulation of GlcNAc-6P and reducing cellular pools of GlcN-6P. Without adequate GlcN-6P levels, *C. rodentium*’s peptidoglycan synthesis pathway was disrupted, compromising cell wall integrity, increasing Δ*nagA* sensitivity to cell wall-targeting antimicrobials as well as osmotic pressure, ultimately resulting in decreased fitness and impairing its ability to establish an *in vivo* infection.

Intestinal mucus is primarily composed of the glycoprotein Muc2. Mucus can function as a physical barrier, preventing luminal microbes from accessing the underlying host tissues. Studies have shown that mice deficient in Muc2 or possessing a thinner than normal mucus layer are more susceptible to enteric infections (16, 34). *In vitro* experiments further demonstrate that mucus inhibits the attachment and translocation of the A/E pathogen EPEC (35–37). However, mucus can also serve as a habitat and provide nutrient sources for certain types of enteric bacteria (38). For example, the mucin-degrading commensal bacterium, *Akkermansia muciniphila*, can modulate the thickness of mucus by secreting mucolytic enzymes, which liberate mucin-derived sugars, including GlcNAc (39). These freed mucin sugars then become accessible carbon sources not only for beneficial commensal microbes but also for enteric pathogens, promoting their expansion and virulence, as shown in *S.* Typhimurium (7), *E. coli* (12, 19, 40, 41), *C. difficile* (7) and *Yersinia enterocolitica* (42).

Building on this broader understanding of mucin sugar utilization, our findings suggest that *C. rodentium* is well-adapted to this nutrient niche. Prior work has shown that *C. rodentium* preferentially colonizes colonic crypts enriched in Muc2-secreting goblet cells (43). Consistent with this, we found that *C. rodentium* can utilize three of the five Muc2-linked sugars as sole carbon sources for growth, with a pronounced preference for GlcNAc and NeuNAc over galactose (Figure 1A). This selectivity might offer a competitive advantage to *C. rodentium* in a nutrient-limited intestinal environment, consistent with Freter’s hypothesis that pathogens must efficiently compete for limited substrates to avoid being flushed out by intestinal flow (44). This is especially relevant for *Enterobacteriaceae* pathogens, including *C. rodentium*, which must traverse or inhabit the mucus layer before infecting the underlying intestinal epithelium (16, 45). To assess the availability of GlcNAc and NeuNAc in the murine intestine, we used lectin staining to demonstrate the spatial distribution (Figure 1B-C). While lectins bind glycan motifs rather than individual monosaccharides-e.g., WGA preferentially binds to terminal GlcNAc residues in dimer and trimer forms (46), and SNA recognizes NeuNAc linked to terminal galactose (47), they remain useful tools for mapping these sugars in colonic tissues. The primary source of free GlcNAc and NeuNAc in the lumen was confirmed to be the Muc2 mucin, as *Muc2^-/-^* mice displayed significantly lower levels of these sugars in their feces, as compared to *Muc2^+/+^*mice (Figure 1D). Moreover, GlcNAc and NeuNAc remain available in the colon throughout *C. rodentium* infection and may serve as accessible nutrients that support *C. rodentium*’s growth and persistence *in vivo* (Figure 1E-F).

Although Δ*mana* alone sustained high colonization levels, its competitive disadvantage against WT in co-infection studies suggests that access to GlcNAc and NeuNAc provides a fitness advantage to *C. rodentium* during infection (Supplementary Figure 1). We selected C57BL/6J mice for this study because they display lower microbiota diversity and reduced colonization resistance against *C. rodentium* when compared to the C57BL/6NCrl used in our prior study with Δ*nanT* (25). The reduced microbial competition in C57BL/6J may obscure the contributions of GlcNAc and NeuNAc due to broader nutrient availability and redundancy in the gut. Supporting this notion, Fabich *et al*. demonstrated that deletion of GlcNAc and NeuNAc transport genes in EHEC EDL933 compromised its fitness only when tested in direct competition with their isogenic parent strains in streptomycin-treated mice (48). These findings underscore the importance of GlcNAc and NeuNAc under competitive conditions, where nutrient utilization becomes a critical determinant.

The Δ*nagA* and Δ*mana* mutants exhibited different colonization outcomes in C57BL/6J mice despite their shared inability to utilize GlcNAc and NeuNAc for growth, with deletion of *nagA* leading to a significantly attenuated phenotype (Figure 2B-E). These findings highlight a potential limitation in interpreting the importance of specific nutrients solely through catabolic gene deletions, as such mutations may also lead to unintended metabolic imbalances or chokepoints. For example, Sinha *et al*. highlighted the importance of GlcNAc for *Enterobacter hormaechei* infection as demonstrated by the reduced colonization of Δ*nagA* in neonatal mice (24). Similarly, Schubert *et al*. described GlcNAc as a context-dependent nutrient source based on the findings that Δ*nagB S.* Typhimurium is less competent than WT *S.* Typhimurium in specific mouse models (23). Our data suggest that it is not the nutrient deprivation *per se* but rather the buildup of metabolic intermediates, namely GlcNAc-6P, that disrupts the normal GlcN-6P synthesis in *C. rodentium* and results in impaired fitness. The Δ*nagA* mutant showed a pronounced growth defect in nutrient-rich LB media, which was further exacerbated following GlcNAc supplementation, suggesting that increased intracellular delivery of GlcNAc was toxic to the cell in the absence of *nagA* (Figure 3B-C). Using the Morgan-Elson assay, we confirmed the accumulation of GlcNAc-6P in Δ*nagA C. rodentium*, establishing a direct link between impaired catabolism and metabolite buildup (Figure 3E). The conservation of the *nagA* gene across *Enterobacteriaceae* suggests that the potential consequences of sugar-phosphate accumulation, particularly GlcNAc-6P, probably occur in an array of enteric pathogens, including *E. hormaechei* and *S.* Typhimurium (Supplementary Figure 2). Our findings highlight the necessity for researchers studying bacterial sugar metabolism to consider the negative impact of metabolite accumulation, which may significantly influence pathogen fitness and virulence, rather than attribute colonization defects solely to nutrient deprivation.

A growth defect resulting from the deletion of *nagA* has been previously reported in *E. coli* (49), *Bacillus subtilis* (50)*, Gluconacetobacter xylinus* (51), and *Streptomyces coelicolor* (52). Unlike common monosaccharides, such as glucose, fructose, and galactose, whose phosphate derivatives have been well described as exerting inhibitory effects on the growth of *E. coli* and *S.* Typhimurium (53), the precise mechanism of GlcNAc-6P toxicity remains poorly understood in Enterobacter species. In many cases, there are two hypothesized causes of sugar-phosphate toxicity. One is that the accumulation of sugar phosphates can directly pose a toxic effect, potentially by forming harmful byproducts (54). A recent study found that GlcNAc-6P can be transformed into a cytotoxic structural analogue of ribose in *Streptomyces* by NagA and a previously uncovered enzyme, NagS, which is not present in *C. rodentium* (55). The other hypothesis is that the buildup of sugar phosphates depletes essential downstream intermediates and eventually impairs energy production or biosynthetic processes (56). Our data support the latter hypothesis because the GlcNAc-6P does not directly kill the Δ*nagA* cells, whereas replenishing the GlcN-6P pool rescues the defect. Unlike in *Streptomyces*, where deletion of *nagA* relieves toxicity by preventing the formation of downstream toxic metabolites (55), *nagA* disruption in *C. rodentium* initiates a stress response that arises from an imbalance in metabolic flux that potentially compromises biosynthetic capacity.

Although Δ*mana* is similarly unable to generate UDP-GlcNAc from exogenous GlcNAc or NeuNAc, it can still readily grow and colonize mice under single infection conditions (Figure 3F-G). This observation implies that it is not merely the absence of GlcNAc or GlcN-6P that overtly impairs fitness but rather the misregulation of GlcN-6P intracellular pools resulting from GlcNAc-6P accumulation. Δ*nagA* cells showed significantly increased expression of *nagB*, indicative of NagC inactivation and enhanced GlcNAc catabolism (Figure 4C). These findings support a model in which the *nagA* deletion not only blocks the breakdown of GlcNAc and NeuNAc into GlcN-6P but also rewires transcriptional regulation to obstruct the synthesis of GlcN-6P from Fru-6P. In contrast, Δ*mana* avoids GlcNAc-6P accumulation and the associated transcriptional misregulation, allowing it to maintain metabolic stability despite being unable to utilize mucin-derived sugars.

This study found that abnormal synthesis of GlcN-6P sensitizes *C. rodentium* to lysozyme, osmotic stress, and cell wall-targeting antimicrobials (Figure 4D-I). These phenotypes underscore the critical role of GlcN-6P in maintaining cell envelope integrity, likely through its involvement in peptidoglycan and lipopolysaccharide biosynthesis. A similar phenomenon has been observed in *Listeria monocytogenes*, where deletion of the *nagA* homolog (*lmo0956*) led to impaired cell wall synthesis, altered cell morphology, reduced teichoic acid content, and heightened sensitivity to mutanolysin, colistin, and ceftriaxone (57). In *E. coli*, NagA is also central to the dedicated peptidoglycan recycling pathway, which salvages GlcNAc released during murein turnover for de novo synthesis of murein and lipopolysaccharide (58). These findings highlight a conserved role for NagA across bacteria families in maintaining a balanced pool of precursors critical for cell wall homeostasis.

Together, these results support a potential therapeutic concept: metabolic sabotage via sugar-phosphate accumulation. By supplying exogenous sugar to the pathogen while simultaneously inhibiting the appropriate enzyme in that sugar utilization pathway like NagA, pathogens may be forced to accumulate toxic intermediates that deplete essential metabolites and compromise their survival. Given that NagA is conserved among *Enterobacteriaceae* (Supplementary Figure 2), it presents a novel target for small-molecule inhibition. For instance, pyrimirhodomyrtone was identified as a natural product that impairs *Staphylococcus aureus* growth by binding the NagA active site (59). Additionally, methyl phosphonamidate analogs mimic the tetrahedral transition state of NagA’s enzymatic reaction and act as tight-binding inhibitors when incubated with NagA purified from *E. coli* (60). While this approach is conceptually attractive, its application *in vivo* would require strategies to selectively target pathogens without disrupting commensals (61).

Future studies should explore identifying pathogen-specific inhibitors, potentially through differences in enzyme structure or regulation, and evaluate their ability to attenuate colonization and virulence. Moreover, combining such intervention with existing cell wall-targeting antibiotics may further enhance therapeutic efficacy.

In summary, our findings show that interruption of GlcNAc and NeuNAc metabolic pathways negatively impacts *C. rodentium* colonization fitness. Upon deletion of *nagA*, GlcNAc-6P accumulation attenuates *C. rodentium* cell wall integrity, sensitizing the bacteria to cell-wall targeting stressors. These findings highlight the role of metabolic context in shaping host-pathogen interactions *in vivo*. Exploiting this vulnerability by targeting key enzymes, such as N-acetylglucosamine-6-phosphate deacetylase, could enable the development of novel therapeutics against A/E pathogens and other enteric microbes.

## Methods

### Ethics statement

All animal experimental protocols were approved by the University of British Columbia’s Animal Care Committee (A23-0204) and all experiments were performed in direct accordance with guidelines drafted by the Canadian Council on Animal Care.

### Bacterial strains and routine growth conditions

All bacterial strains used in this study are listed in Table S1. Bacteria were routinely cultured in Luria Broth (LB) at 37°C aerobically with shaking at 200 rpm. LB supplemented with 1.5% agar was used for solid medium growth with aerobic incubation at 37°C for 16-18 hours. When required, the growth medium was supplemented with streptomycin (Catalog #S4014, Sigma-Aldrich) at 100 µg/mL.

### In vitro growth assays

To assess the ability of *C. rodentium* to utilize and grow on Muc2-associated monosaccharides, M9 minimal media was used and generated by first preparing a 10X autoclaved base media (37.07 g of Na_2_HPO_4_, 30 g of KH_2_PO_4_, 5 g of NaCl, and 10 g of NH_4_Cl per a litre), then diluted in ddH_2_O to a final concentration of 2 mM MgSO_4_, 0.1 mM CaCl_2_, and 0.1% w/v of the sugar source. The final concentration of sugars was selected based on the reported concentration of these sugars in the purified porcine gastric mucin and to mimic the nutrient-limited conditions in the intestinal environment (62, 63). Sugar sources include GlcNAc, NeuNAc, GalNAc, galactose, and fucose. To evaluate the susceptibility of *C. rodentium* to antibiotics, LB was used as the growth medium supplemented with 64 µg/mL vancomycin or 8 µg/mL tetracycline, which is half of the minimal inhibitory concentration for WT *C. rodentium*. To monitor the growth of *C. rodentium,* 200 µL of the growth media was added to each well of the clear-bottom 96-well plate. Overnight cultures of *C. rodentium* grown in LB were washed with PBS and added to the media with the optical density measured at 600 nm (OD_600_) starting at 0.01. The cultures were incubated in the automated VarioskanTM LUX multimode plate reader (Thermo Fisher, USA) at 37 °C with orbital shaking at 180 rpm. The growth was monitored by the spectrophotometer at OD_600_ for 18 h, and data were analyzed on Skanlt software (version 6.1).

### Measurements of Muc2 sugars using UHPLC/QqQ-MS

Quantification of GlcNAc and NeuNAc within mouse feces was adapted from Xu *et al*. (64), with slight modifications in the extraction, derivatization, and LC-MS conditions. Briefly, fecal samples (∼20–30 mg) were diluted 20-fold (final concentration 20 µL/mg) with distilled water and homogenized overnight at 4 °C. Supernatants were clarified by centrifugation (21,000 ×g, 20 min) and subjected to 1-phenyl-3-methyl-5-pyrazolone (PMP) derivatization. Standard curves were prepared from a mixture of GlcNAc, galactose, and glucose, while NeuNAc standards were prepared separately and added after derivatization (0.39–50 µg/mL). For derivatization, 50 µL of sample or standard was mixed with 200 µL of 20–22% ammonia and 200 µL of 0.2 M PMP in methanol, incubated at 70 °C for 30 min. NeuNAc (not derivatized) was added afterward. Final mixtures of 20 µL derivatized sample and 60ul non-derivatized sample were diluted 25-fold with dH₂O and 10% acetonitrile before injection. Quantitation of the sample sugars was done using an external standard method.

UHPLC separation was performed on a Waters Xevo TQS triple quadrupole mass spectrometer equipped with a Waters H-class Acquity UHPLC system and a Waters Acquity UHPLC BEH C18 Waters Acquity BEH C18 column (2.1 × 100 mm, 1.7 µm) at 35 °C with a flow rate of 0.4 mL/min. The aqueous mobile phase A was 5 mM ammonium acetate adjusted to pH 8.3 with ammonium hydroxide; mobile phase B consisted of 95% acetonitrile and 5% ammonium acetate buffer (v/v, pH 8.3). The optimized gradient was as follows: 0.0–7.0 min, 12–15% B; 7.1–8.5 min, 99% B; 8.6–10.0 min, 12% B. All four compounds were baseline resolved, with retention times of ∼0.5 min (NeuNAc), ∼5.0 min (Glc), ∼5.5 min (galactose), and ∼6.5 min (GlcNAc). Detection was performed on the MS in MRM mode. Transitions were 511.2→175 (glucose/galactose), 552.2→175 (GlcNAc), and 310.3→274.2 (NeuNAc), with respective collision energies of 25, 30, and 5 eV. All analyses were performed using MassLynx and QuanLynx software.

### Construction of C. rodentium mutants and complementation strains

In-frame deletion mutants of *C. rodentium* strain DBS100 (streptomycin-resistant) were constructed using overlap extension PCR (65) and allelic exchange (66) strategies as previously described (67). Briefly, genomic regions (∼1kb) flanking the target gene were amplified using primers listed in Table S2 with overlapping sequences for fusion (P2 and P3) and engineered restriction sites (P1 and P4). The two fragments were fused and cloned into the suicide vector pRE112, which confers chloramphenicol resistance for selection. The resulting construct was transformed into *E. coli* MFD λpir by electroporation and conjugated into the WT *C. rodentium* strain. Mutants were selected based on sucrose resistance and loss of chloramphenicol resistance, indicative of successful double crossover events, and were confirmed by PCR and Sanger sequencing using gene-specific check primers (PF and PR).

Genetic complementation was performed as previously described (68). Specific primers (Table S2) were used to amplify the native promoter (P1-comp and P2-comp) and coding sequence of *nagA* (P3-comp and P4-comp). The construct was cloned into the XhoI/BamHI sites of pZA31MCS (Expressys, Ruelzheim, Germany) (69) to generate the plasmid pNagA containing wild-type *nagA*. Plasmid pNagA was electroporated into the *C. rodentium* Δ*nagA* strain to produce the *C. rodentium* Δ*nagA*/pNagA complemented strain.

### In vivo mice infection studies

C57BL/6J (6 to 8 weeks old) female mice were obtained from the Jackson Laboratory and maintained at the BC Children’s Hospital Research (BCCHR) Institute. Mice were kept in sterilized cages with filter tops, handled in tissue culture hoods, and provided with autoclaved food and water under specific pathogen-free conditions. Sentinel animals were routinely tested for common pathogens. For infection, mice were orally gavaged with 0.1 ml of overnight LB culture (∼ 2.5x10^8^ CFU) of *C. rodentium*. To monitor colonization, fecal samples were collected in PBS, at specified time points, weighed, homogenized, serial diluted and plated on streptomycin-containing LB agar plates. The CFU was counted and divided by stool weight.

For tissue collection, mice were euthanized at 8 DPI, and their large intestines were collected and divided into ceca, proximal, and distal colons. For histological studies, a portion of the ceca and distal colons was obtained and fixed in either 10% formalin or Methacarn solution (methanol: chloroform: acetic acid, 6:3:1, v/v), where tissue sections were obtained with retained luminal content to preserve the mucus architecture. Next, the remainder of the ceca and distal colons were opened to retrieve the luminal content and thoroughly washed in PBS. Tissue and luminal contents were placed in Eppendorf tubes with 1 mL PBS and beads for bacterial enumeration, similar to the stool samples described above.

### In vivo competitive assays

The *C. rodentium* WT-AC strain and the competitive assay were conducted with modifications for *in vivo* infection following the protocol previously described by Gilliland et al (67). In brief, the WT-AC *C. rodentium* expressing a tetracycline-inducible amCyan fluorescent protein was used to compete with Δ*mana*. The overnight culture of the two strains was normalized to the same OD_600_ and mixed in a 1:1 ratio. 100 µL of the mixture was orally gavaged to mice. Stool samples were collected in the following days, serially diluted, and plated on streptomycin-containing LB agar plates with 0.2 μg/mL anhydrotetracycline (Fisher, Catalog #AAJ66688MA) to induce the expression of amCyan in WT-AC. Plates were imaged using the iBright FL1500 Imaging System (Thermo-Fisher USA) for visualization of fluorescent WT-AC. The number of Δ*mana* (nonfluorescent) colonies was determined by subtracting the number of fluorescent colonies from the total colony count. The number of colonies corresponding to WT-AC versus Δ*mana C. rodentium* was transformed into a fraction of 1 to calculate the competitive index value.

### Histology and scoring

For routine histology to determine tissue morphology and pathophysiology changes, cecal and colonic tissues were fixed in 10% formalin, paraffin-embedded, and sectioned at 5 µm and then stained with hematoxylin and eosin (H&E). Histopathological scores were determined by two blinded observers independently as previously described (70). In brief, readouts included submucosal edema (0, no edema; 3, profound edema), epithelial hyperplasia (score based on the percent change in crypt height compared to that of the control crypts; 0, no change; 1, 1% to 50% change; 2, 51% to 100%; 3, >100%), polymorphonuclear cell infiltration (0, none; 3, severe), and epithelial integrity (0, no damage; 1, 10 epithelial cells shedding per lesion; 2, 11 to 20 epithelial cells shedding per lesion; 3, maximum damage to epithelial surface as noted by crypt destruction and epithelial ulceration). The maximum possible score was 12.

### Lectin staining

Paraffin-embedded colonic tissue sections (5 μm) were deparaffinized by heating at 60 °C for 15 min, cleared with xylene, and rehydrated with 100%, 95%, and 70% ethanol, followed by dH_2_O. Dewaxed and dehydrated colonic tissue sections were blocked with Donkey Serum buffer at room temperature (RT) for 1 hour. Fluorescently labelled lectins were diluted in antibody dilution buffer and used for staining: Wheat Germ Agglutinin (WGA) (Catalog #FL-1021, Vector Laboratories, 1:500) for GlcNAc, and Sambucus Nigra Lectin (SNA) (Catalog #FL-1301, Vector Laboratories, 1:200) for NeuNAc. For negative controls, lectins were pre-incubated with 100 mM of their target sugars for at least 30 min in the dark, then applied to unstained sections. All staining was carried out at RT for 2 hours. Slides were washed once in PBS, twice in PBS + 0.1% TritonX-100, and twice in PBS. Sections were then mounted using ProLong Gold Antifade reagent (Molecular Probes/Invitrogen) that contains 4′,6′-diamidino-2-phenylindole (DAPI) for DNA staining. Sections were viewed at 350, 488, and 594 nm on a Zeiss AxioImager microscope. Images were obtained using a Zeiss AxioImager microscope equipped with an AxioCam HRm camera operating through AxioVision software (Version 4.4).

### Intracellular GlcNAc(6P) quantification

The level of GlcNAc(6P) was estimated by a modification of the Morgan-Elson procedure improved by Plumbridge (71). Briefly, bacteria were grown in 25 mL minimal media supplemented with 0.1% Glc and GlcNAc for 24 hours. Each sample was normalized to OD_600_ = 0.5 (10^7^∼10^8^ CFU/mL) and then harvested by centrifugation at 4200 rpm, 4°C for 10 min. Pellets were washed with 1.5 mL H_2_O, resuspended in 250 µL H_2_O, and placed in a boiling water bath for 5 min for metabolite extraction. 50 µL bacterial extracts were mixed with 75 µL of 2 M potassium tetraborate tetrahydrate (Catalog #P5754, Sigma-Aldrich). The extracts were boiled for 3 min, cooled to room temperature, and then incubated with 625 µL Ehrlich’s reagent, prepared by dissolving 1 g of 4-(dimethylamino) benzaldehyde (Catalog #109762, Sigma-Aldrich) in 1.25 mL HCl and 100 mL glacial acetic acid, for 25 min at 37°C. The reacted purple extracts were centrifuged at 13000 rpm at 4 °C for 5 min to cease the reaction and remove any precipitate. The absorbance of the extracts was measured at OD_585_. GlcNAc standards (125 – 2000 µg/mL) were used to generate a standard curve for calculation.

### Lysozyme killing assay

Overnight cultures of *C. rodentium* were diluted in PBS to an OD_600_ of 0.01 (∼10^6^ CFU/mL). The bacteria culture was then treated with 10 mg/mL lysozyme (Catalog #L6876, Sigma-Aldrich), with or without supplementation of 0.1% GlcNAc and GlcN as indicated. Samples were incubated for 18 h at 37°C. After incubation, samples were serially diluted and plated on streptomycin-containing LB agar plates for CFU enumeration. Relative viability was calculated by N/N_L_, where N and N_L_ represent CFU counts in the presence or absence of lysozyme, respectively.

### Osmolarity viability assay

Overnight cultures of *C. rodentium* were diluted into 3 mL of LB broth with either 1% or 3% NaCl to an initial OD_600_ of 0.01. After 3 h of incubation at 37°C, bacterial suspensions were serially diluted and plated on LB agar to determine CFUs. Relative viability was calculated by N_3_/N_1_, where N_3_ and N_1_ represent CFU counts in the 3% and 1% NaCl conditions, respectively.

### RNA extraction and quantitative PCR

Overnight cultures of *C. rodentium* were diluted and incubated in 7 mL LB broth at 37°C starting at OD_600_ = 0.01, and were harvested when OD_600_ = 0.7. Total RNA was isolated using the RNeasy Mini kit (Qiagen) according to the manufacturer’s protocol, and the RNA concentration was measured at the NanoDrop spectrophotometer (Thermo Fisher). Extracted RNA was reverse transcribed using 5X All-In-One RT MasterMix (Applied Biological Materials). For quantitative real-time PCR (qPCR), the cDNA was diluted 1:5 in RNase/DNase-free water. 5 μl of the diluted cDNA was added to the PCR mix containing 10 μl of SsoFast EvaGreen Supermix (Bio-Rad) and primers listed in Table S3 (300 nM). qPCR was performed on a Bio-Rad CFX connect Real-time PCR detection system with the following cycling conditions: denaturation at 95°C for 30 s, followed by 3 s of denaturation at 95°C, 5 s of annealing at 60°C, 5 s of extension at 65°C for a total of 39 cycles. mRNA transcript expression was normalized to the relative expression of the reference gene, *rpoA*, using the 2^-(ΔCt)^.

### Phylogenetic and genomic analyses

A phylogenetic tree of *Enterobacteriaceae* bacteria containing *nagA* (K01204; EC 3.2.1.49) was generated using AnnoTree v1.2 at the genus level. Searches for *nagA* were performed with a minimum of 95% identity, 95% subject alignment, 95% query alignment, and an E value cutoff of 1 × 10⁻⁵. This resulted in the selection of the following genera: Yersinia, Vibrio, Shewanella, Serratia, Salmonella, Proteus, Pasteurella, Pantoea, Klebsiella, Hafnia, Haemophilus, Escherichia, Erwinia, Enterobacter, Cronobacter, Citrobacter A, Citrobacter, and Aeromonas. In R (v4.5.0), duplicate species entries were removed, and for each genus, the total number of genomes and the number and percentage of genomes containing *nagA* were calculated. Genera were categorized by total genome counts (0–10, 11–50, and >50). A heatmap of the percentage of genomes containing *nagA* was generated using the pheatmap package (v.1.0.13), with genome count categories displayed as row annotations and percentage values shown directly on the heatmap.

### Statistical analyses

Statistical analyses were performed using GraphPad Prism version 9.5.0 (GraphPad Software Inc.). Differences between two groups were evaluated using either the Student’s *t*-test (parametric) or the Mann-Whitney U test (non-parametric), with the test that was applied noted in each figure legend. For multiple comparisons, statistical analysis was performed using one-way ANOVA (parametric) or Kruskal-Wallis test (non-parametric), and then the Dunnett’s or Bonferroni test for parametric samples or Dunn’s test for non-parametric samples as a post-hoc test. Differences at p < 0.05 were considered significant, with asterisks denoting significance in figures.

## Acknowledgement

We greatly appreciate the technical support from Caixia Ma and the BCCHRI animal staff for the mouse experiment. We also acknowledge assistance from Ashley Gilliland in mutant engineering and Roger Dyer in mucin sugar quantification. We greatly thank Yuchan Zhu for her support in editing immunofluorescent and H&E images. This study was funded by the Canadian Institutes of Health Research (http://www.cihr-irsc.gc.ca/e/193.html) AWD-019071 and AWD-026220, as well as the Natural Sciences and Engineering Research Council of Canada, to B.A.V. B.A.V. is the Children with Intestinal and Liver Disorders (CH.I.L.D.) Foundation Chair in Pediatric Gastroenterology.

## Author contributions

Zhiquan Clarence Huang, Conceptualization, Methodology, Validation, Formal analysis, Investigation, Writing – original draft, Writing – review and editing, Project administration | Matthias Ahmad Mslati, Conceptualization, Formal analysis, Investigation, Data curation, Investigation, Project administration | Caixia Ma, Methodology, Resources, Data curation | Hyunjung Yang, Validation, Writing – original draft, Writing – review and editing, Supervision | Qiaochu Liang, Data curation, Resources, Writing – original draft, Writing – review and editing, Supervision | Shauna Crowley, Visualization, Writing – original draft, Writing – review and editing | Roger Dyer, Methodology, Formal analysis, Data curation | Irvin Ng, Software, Formal analysis | Hongbing Yu, Conceptualization, Writing – original draft, Writing – review and editing, Project administration, Supervision | Bruce A. Vallance, Conceptualization, Resources, Writing – original draft, Writing – review and editing, Project administration, Supervision, Funding acquisition

### Data availability

All data are contained within the manuscript.

